# Chronic exposure to a neonicotinoid pesticide and a synthetic pyrethroid in full-sized honey bee colonies

**DOI:** 10.1101/293167

**Authors:** Richard Odemer, Peter Rosenkranz

## Abstract

In the last decade, the use of neonicotinoid insecticides increased significantly in the agricultural landscape and meanwhile considered a risk to honey bees. Besides the exposure to pesticides, colonies are treated frequently with various acaricides that beekeepers are forced to use against the parasitic mite *Varroa destructor*. Here we have analyzed the impact of a chronic exposure to sublethal concentrations of the common neonicotinoid thiacloprid (T) and the widely used acaricide τ-fluvalinate (synthetic pyrethroid, F) - applied alone or in combination - to honey bee colonies under field conditions. The population dynamics of bees and brood were assessed in all colonies according to the Liebefeld method. Four groups (T, F, F+T, control) with 8-9 colonies each were analyzed in two independent replications, each lasting from spring/summer until spring of the consecutive year. In late autumn, all colonies were treated with oxalic acid against Varroosis. We could not find a negative impact of the chronic neonicotinoid exposure on the population dynamics or overwintering success of the colonies, irrespective of whether applied alone or in combination with τ-fluvalinate. This is in contrast to some results obtained from individually treated bees under laboratory conditions and confirms again an effective buffering capacity of the honey bee colony as a superorganism. Yet, the underlying mechanisms for this social resilience remain to be fully understood.

## 1 INTRODUCTION

Neonicotinoid pesticides are among the most used insecticides during the past decades and are dominating the global market for insecticidal seed dressings (Jeschke et al., 2011; Simon-Delso et al., 2015). However, these neonicotinoids are suspected to be a main driver for the decline of honey bees (Hopwood et al., 2016), wild bees (Potts et al., 2010) and even non-target wildlife in general (Goulson, 2013). Recently, the European Food Safety Authority (EFSA) has updated their risk assessment and now considers the three neonicotinoids imidacloprid, clothianidin and thiametoxam to be “a risk for bees” and suggested suitable amendments to the European Commission (EFSA, 2018). These three nitro-substituted compounds have the highest toxicity to bees among the class of neonicotinoids (Iwasa et al., 2004) and have been already banned for the use in flowering crops by the European Union since the year 2014 (EFSA, 2013).

However, other neonicotinoid insecticides with a far lower toxicity to bees - for instance thiacloprid and acetamiprid - are still widely used not only as seed dressings but are even approved as foliar spray in blooming cultures like oilseed rape (Schmuck et al., 2003). This leads to a remarkable high contamination of nectar and pollen and foragers might therefore be continuously exposed to these agents (Genersch et al., 2010; Collison et al., 2016; Rolke et al., 2016; Böhme et al., 2017). There is no doubt about the comparable low acute toxicity of these compounds to bees, however there is a controversial discussion on sublethal and long-term effects. So, it has been shown that thiacloprid can affect the sensitivity of honey bees to the gut parasite *Nosema ceranae* (Vidau et al., 2011; Pettis et al., 2013; Retschnig et al., 2015). More recent publications indicate that sublethal concentrations of thiacloprid alter their social behavior (Forfert and Moritz 2017) and, more importantly, disturb the orientation of foragers (Fischer et al., 2014; Tison et al., 2016, 2017). These studies have been conducted on the level of individual or small groups of bees by performing cage tests or semi-field trials under rather artificial conditions. Therefore, they do not cover important attributes of a social entity, with a more complex perception to its environment. Hence, the transfer of these results to field conditions must be taken with caution. Significantly, the only field study available so far could not confirm negative effects of thiacloprid at the colony level (Siede et al., 2017).

Another controversial point is the possible interaction of thiacloprid - considered as “non-toxic for bees” - with active compounds of other chemical classes that are applied by beekeepers to control the parasitic mite *Varroa destructor*, requiring multiple annual treatments (Rosenkranz et al., 2010). In an effective and easy to use application, synthetic pyrethroids were, amongst others, introduced to beekeepers (Watkins, 1997) and are besides the formamidine amitraz the most frequently used acaricides in apiculture (Garrido et al., 2016). The exposure of honey bee colonies to a combination of sublethal doses of such pesticides may increase the susceptibility to pathogens and are suspected to contribute to the worldwide health problems of honey bee colonies (Cornman et al., 2013; Matsumoto, 2013; Wu et al., 2012). To study such possible combination effects we have chronically exposed full-sized colonies to the neonicotinoid thiacloprid and the synthetic pyrethroid τ-fluvalinate (Apistan^®^) in a two-year field study. To our knowledge this is the first study that analyzes the effect of a chronic application of both, a neonicotinoid insecticide and a common acaricide under realistic field conditions at the colony level. An exposure to these two pesticides is very likely under common beekeeping conditions in rural areas. Our crucial endpoints were (i) the overwintering success of treated colonies compared to untreated controls and (ii) the colony population dynamics.

## 2 MATERIALS & METHODS

### 2.1 Experimental colonies

For each treatment group, five experimental colonies were established in early May of the year 2010. The experiment was repeated with three to four new colonies per group in the year 2011 (Tab. 1). All colonies were set up at our local apiary at the agricultural experimental station Kleinhohenheim, which is an organic farming facility not using any agro chemicals or common pesticides at all. To standardize our experiment, we used artificial swarms made from stock colonies that were screened for low *Varroa* infestation and lack of virus infections prior to the trials. Freshly reared and mated sister queens of the Hohenheim breeding line were provided to each swarm, respectively. After the colonies successfully showed the first open brood stages, we sprayed all of them with a 3.5 % oxalic acid sugar solution for *Varroa* treatment to have a comparable low mite infestation for all experimental groups at the start of the experiment. We used residue free beeswax foundations to minimize the risk of additional contamination through pesticide residues in the wax (Bogdanov et al., 1998; Wallner, 1999). All colonies were set up on one box of 10 Zander frames, which was extended to two boxes when necessary during the summer season.

**Tab. 1.** List of replications, treatment groups, treatment duration, assessment dates (AD) and no. of colonies (N) at the time of the assessment.

| Year | Treatment | Duration [days] | AD 1) | N | AD 2) | N | AD 3) | N | Winter treatment | N | AD 4) | N |
| --- | --- | --- | --- | --- | --- | --- | --- | --- | --- | --- | --- | --- |
| 2010-2011 | Control | 56 | 23. Jul | 5 | 16. Aug | 5 | 8. Oct | 5 | 30. Nov | 4 | 15. Apr | 4 |
|  | Thiacloprid |  |  | 5 |  | 5 |  | 5 |  | 3 |  | 3 |
|  | Fluvalinate |  |  | 5 |  | 5 |  | 5 |  | 5 |  | 5 |
|  | Flu + Thia |  |  | 5 |  | 5 |  | 5 |  | 4 |  | 4 |
| 2011-2012 | Control | 62 | 21. Apr | 3 | 5. Aug | 3 | 13. Oct | 3 | 29. Dec | 3 | 3. Apr | 2 |
|  | Thiacloprid |  |  | 4 |  | 4 |  | 4 |  | 4 |  | 4 |
|  | Fluvalinate |  |  | 3 |  | 3 |  | 3 |  | 3 |  | 3 |
|  | Flu + Thia |  |  | 3 |  | 3 |  | 3 |  | 3 |  | 3 |

### 2.2 Thiacloprid application

For the application of thiacloprid we used the pure substance (98 % purity, Dr. Ehrenstorfer GmbH), which was sonicated in pure water for a stock solution. We aimed to use a field-realistic concentration that was approximately 100-fold lower than the oral LD_50_ for thiacloprid (173.2 mg/kg, Würfel, 2008). We therefore diluted thiacloprid in sucrose syrup (Apiinvert, Südzucker GmbH) in order to receive the respective concentration. The final solution was quantified by an external lab (Eurofins Dr. Specht Laboratorien GmbH, Hamburg, Germany) which confirmed a thiacloprid concentration of 1.6 mg/kg (= 1,600 ppb). This feeding solution was applied to the colonies of the specific treatment groups and control colonies were fed with untreated sucrose syrup. The duration of the treatment in the year 2010 was 56 days (23^rd^ Jul-17^th^ Sep) and in the year 2011 62 days (21^st^ Apr-22^nd^ Jun) during summer season. In this time period we fed 1 kg syrup per week with an internal feeding device, to simulate a chronic exposure. A final amount of 8 kg per colony in 2010 and 9 kg in 2011 was administered in the summer season, respectively. Based on the concentration of 1.6 mg/kg we therefore applied a total amount of 12.8 mg thiacloprid per colony in 8 weeks (2010) and 14.4 mg thiacloprid per colony in 9 weeks (2011) during the summer season, respectively. The treatment was resumed when colonies were fed for overwintering at the end of the season. Every colony was fed with approximately 15 kg of the feeding solution with a total amount of 24.0 mg thiacloprid in each year for winter feeding. After the treatment period in summer, a pooled sample of food (nectar/honey) from the combs was analyzed for residues at Eurofins Dr. Specht Laboratorien GmbH.

### 2.3 τ-fluvalinate application

Apistan^®^ strips (Vita Europe Ltd, Basingstoke, UK) were used for the τ-fluvalinate treatment. As recommended, one strip per box was applied to the τ-fluvalinate treatment groups during the same time of the thiacloprid application. After the treatment period, a pooled sample of beeswax was analyzed for residues at our own lab in Hohenheim. During overwintering, the strips were again inserted to the colonies to resume a chronic treatment.

### 2.4 Assessment of population dynamics

The amount of bees and brood cells (open and sealed) were estimated with the Liebefelder Method (Imdorf et al., 1987), which is a feasible tool that provides accurate and reliable results at the colony level (measuring error +/-10 %). Care was taken that all colonies were evaluated by the same person on all dates to minimize variation. Colony assessments were usually conducted in the morning before bee flight.

### 2.5 *Varroa* winter treatment

In order to monitor the level of mite infestation in the colonies and to measure the effectiveness of the τ-fluvalinate treatment, we applied 3.5 % oxalic acid sugar solution to the bees in a brood free stage during late autumn or winter time (30^th^ Nov in 2010 and 29^th^ Dec in 2011). In both years the temperature was below 3 °C for optimal application to a closely spaced bee cluster. Dead mites were counted approximately one week after the treatment with a sticky board, which was inserted at the same day of treatment, respectively.

### 2.6 Statistical analysis

The estimated number of bees and brood cells from both years were checked with a Shapiro-Wilk test for normal distribution (p>0.05). Therefore, a one-way ANOVA and a multiple comparison of the means with a post-hoc Bonferroni correction were performed on the four experimental groups, respectively (α=0.05).

All tests were performed using WinSTAT (R. Fitch Software, Bad Krozingen).

## 3 RESULTS

### 3.1 Overwintering success

In both years, none of the colonies died until the start of wintering in October (Tab. 1). Taken both years together, a total of five of the 33 colonies died over winter. Two of the “Thiacloprid” group (N = 9), one of the “Flu+Thia” group (N = 8), two of the “Control” group (N = 8) and none of the “Fluvalinate” group (N = 8; Tab. 1).

### 3.2 Population dynamics

#### 3.2.1 Experiment 1 (2010 - 2011)

The population of bees and brood cells were estimated four times during the whole season (Tab. 1). The results are shown in Fig. 1a for the number of bees and in Fig. 1b for the number of brood cells. We compared the four treatment groups for each date of the estimates and could not see significant differences (ANOVA) for the number of bees in August 2010 (“AUG”; p=0.254), October 2010 (“OCT”; p=0.473) and April 2011 (“APR”; p=0.388). Likewise, no significant differences of the amount of brood cells were recorded in October 2010 (“OCT”; p=0.590) and April 2011 (“APR”; p=0.128). However, in July the number of bees of the “Control” were significantly lower compared to “Fluvalinate” (p=0.029, ANOVA). The number of brood cells of the “Control” was significantly lower compared to “Thiacloprid” and “Flu+Thia” in July (p=0.012, ANOVA) and compared to “Thiacloprid” in August (p=0.004, ANOVA).

**Fig. 1a.**
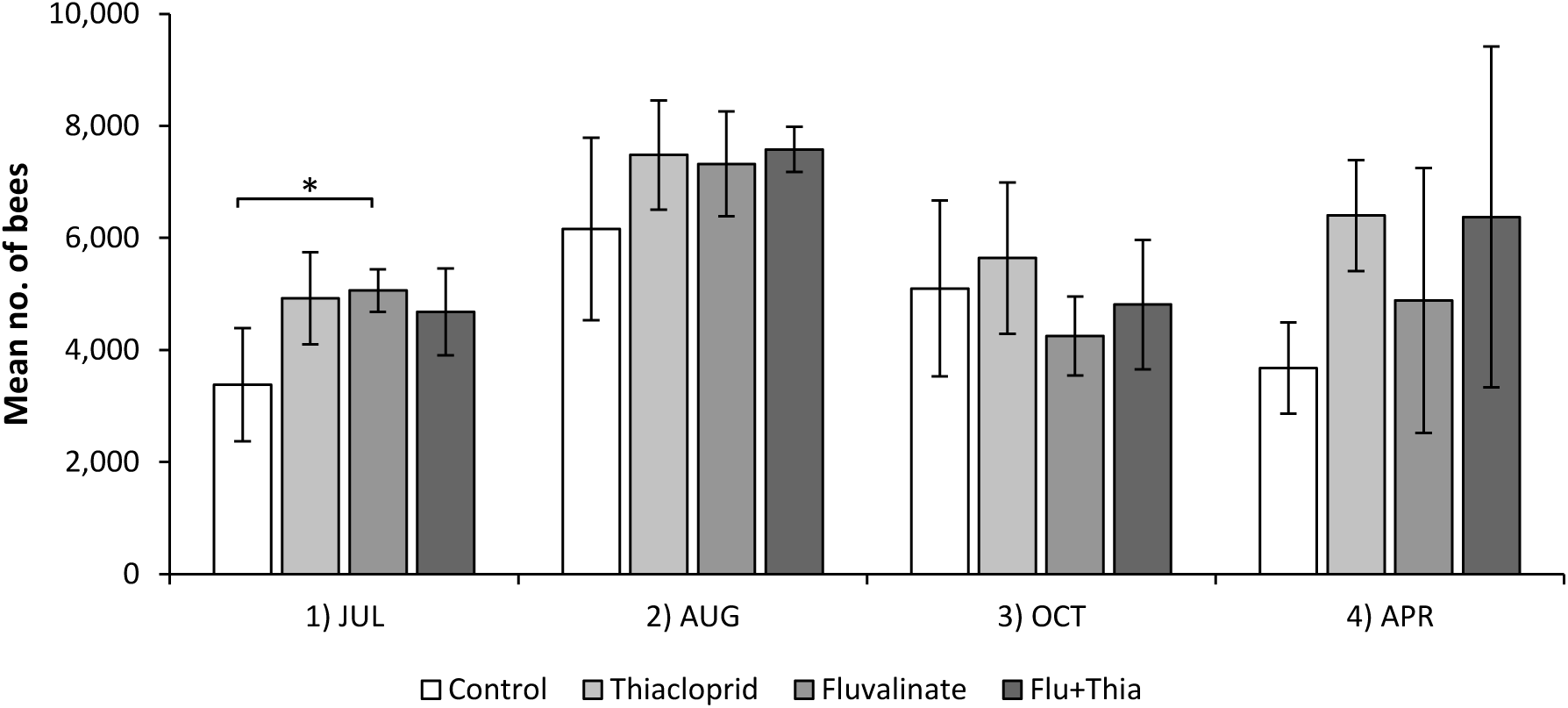
Number of bees estimated in the colonies in the year 2010-2011 for the four treatment groups at four different assessments. * statistically significantly lower when “Control” compared to “Fluvalinate” (p<0.05, ANOVA).

**Fig. 1b.**
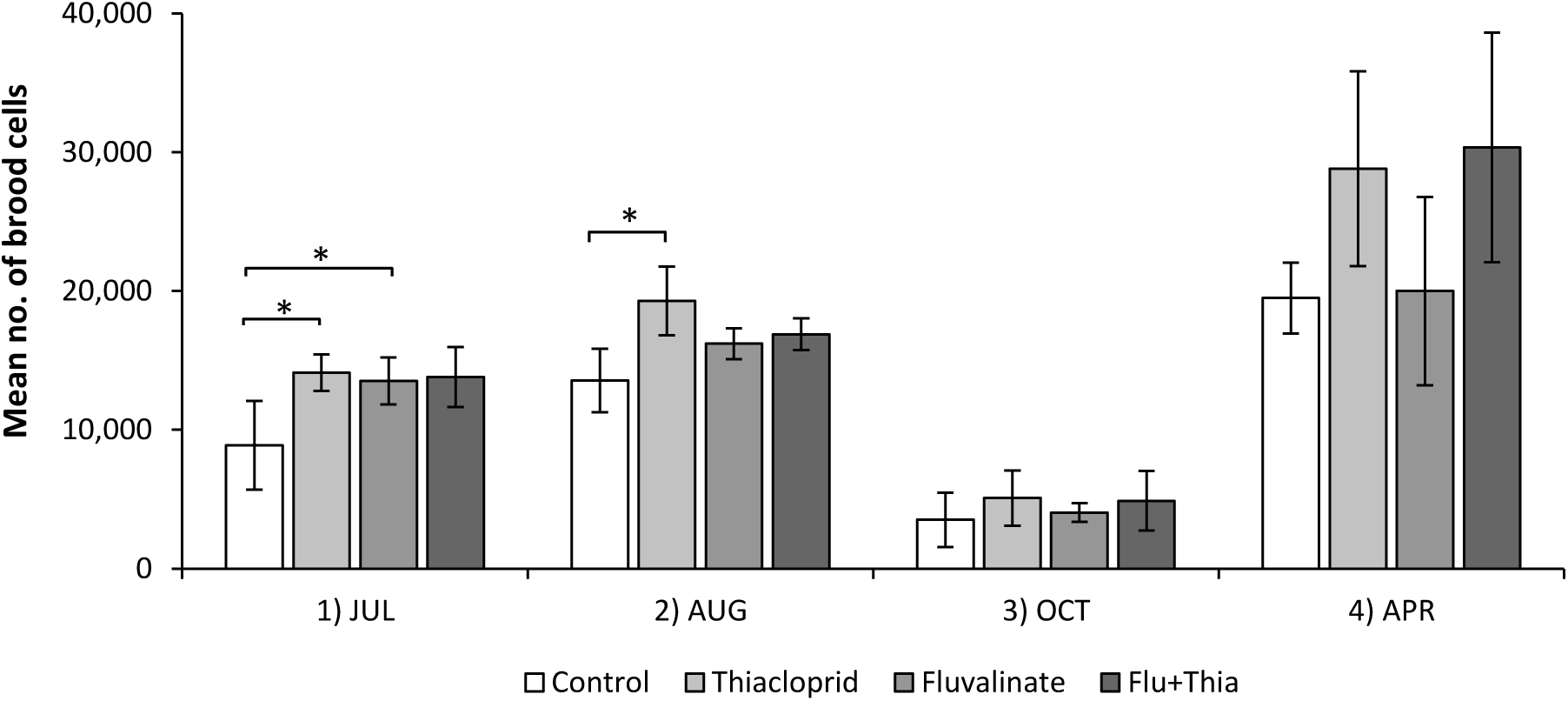
Number of brood cells estimated in the colonies in the year 2010-2011 for the four treatment groups at four different assessments. * statistically significantly lower when “Control” compared to “Thiacloprid” and “Fluvalinate” (p<0.05, ANOVA) in 1), and when “Control” compared to “Thiacloprid” (p<0.05, ANOVA) in 2).

#### 3.2.2 Experiment 2 (2011 - 2012)

For the replicate of experiment 1, also four assessments were performed throughout the season. The results are shown in Fig. 2a for bees and in Fig. 2b for brood. We again compared the four groups within each assessment but could not see any significant differences for the number of bees (April 2011 p=0.174; August 2011 p=0.367; October 2011 p=0.664; April 2012 p=0.198) and no significant differences for the number of brood cells in April 2011 (p=0.071), October 2011 (p=0.328) and April 2012 (p=0.176; ANOVA). Solely, in August 2011, the number of brood cells in “Thiacloprid” was significantly lower compared to “Control” and “Fluvalinate” (p=0.017, ANOVA).

**Fig. 2a.**
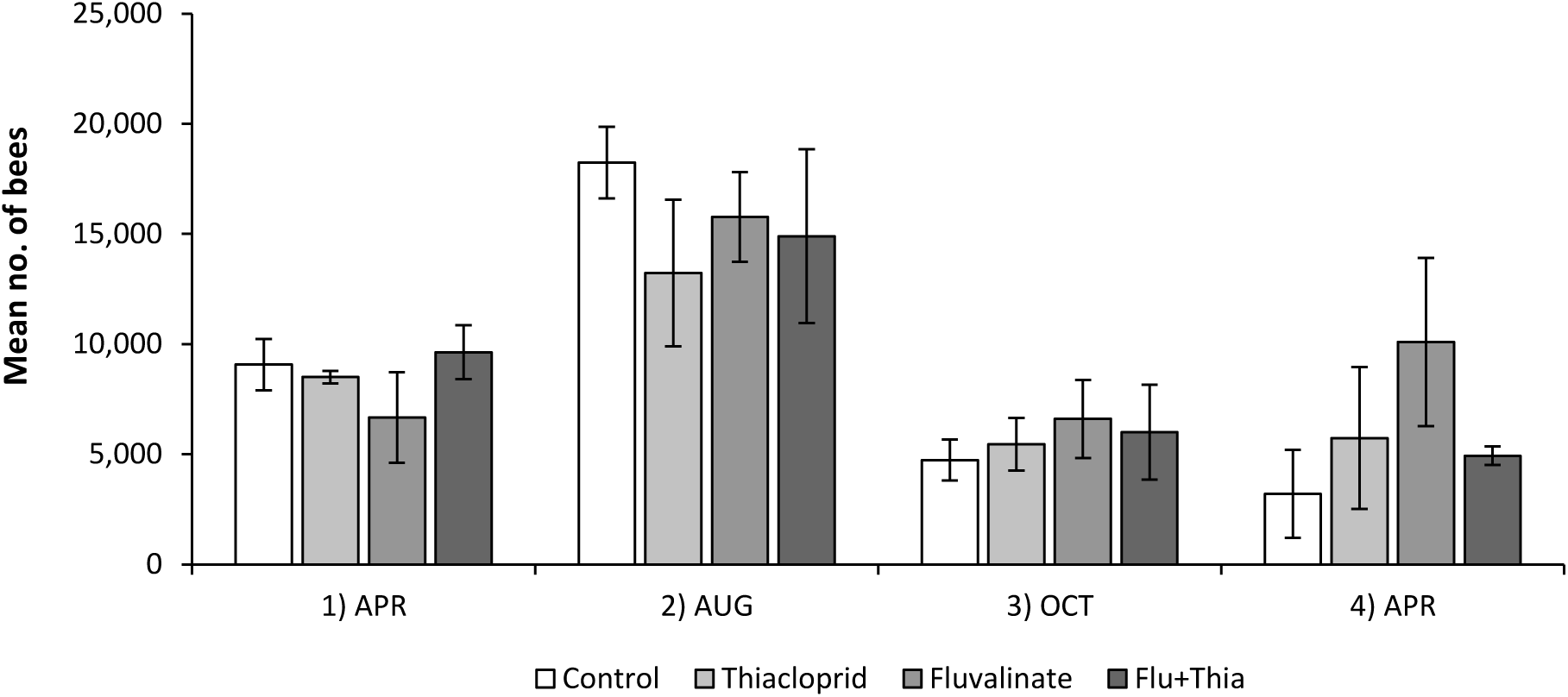
Number of bees estimated in the colonies in the year 2011-2012 for the four treatment groups at four different assessments. We could not see statistically significant differences within the assessments (p>0.05, ANOVA).

**Fig. 2b.**
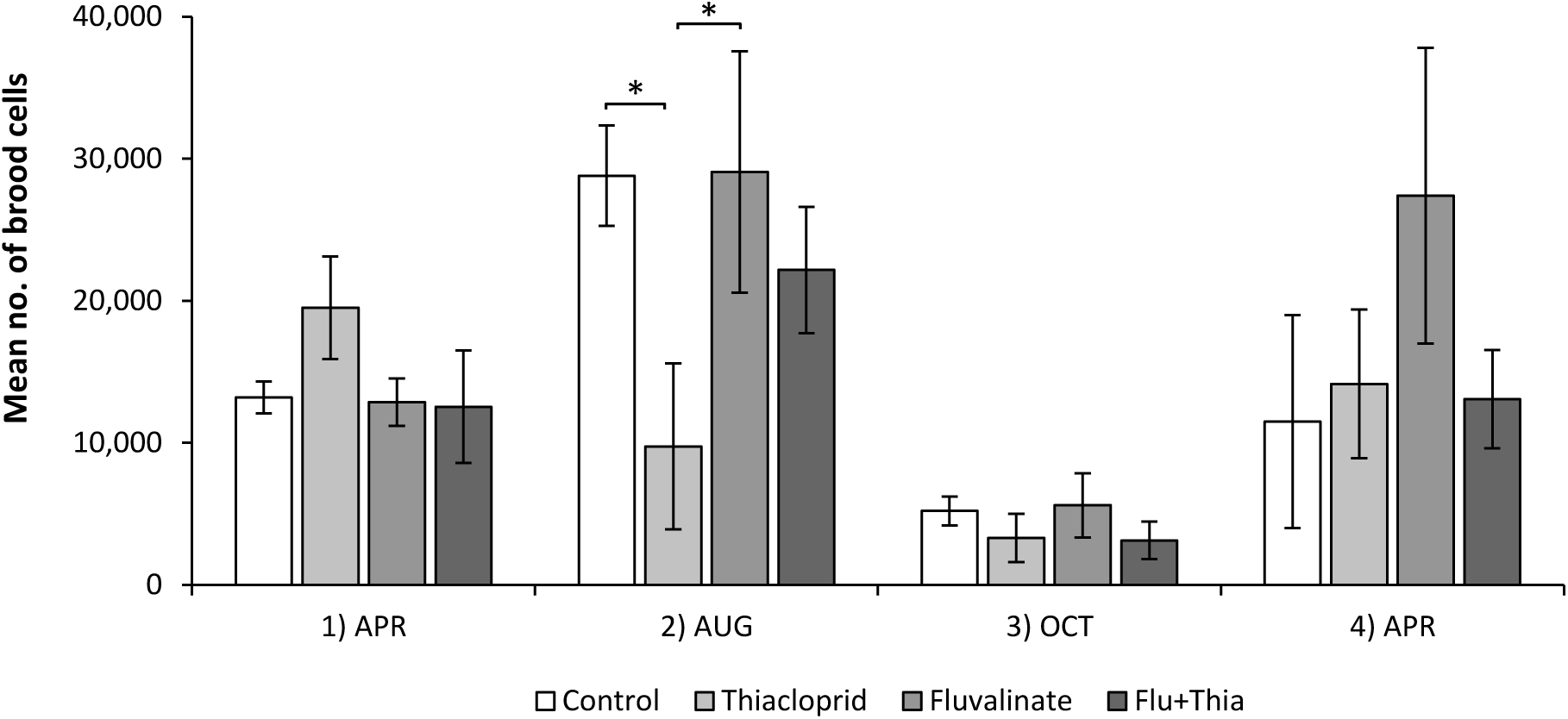
Number of brood cells estimated in the colonies in the year 2011-2012 for the four treatment groups at four different assessments. * statistically significantly lower when “Thiacloprid” compared to “Control” and “Fluvalinate” (p<0.05, ANOVA) in 2).

### 3.3 Thiacloprid residues

Food from the syrup feeding, which was processed by the bees and stored in honeycombs, was analyzed for thiacloprid residues in both years with QuEChERS method (Limit of Quantification LOQ = 0.01 mg/kg). For the analysis, samples from all colonies and the respective groups per year were pooled. All groups without thiacloprid treatment did not have measurable residues in both years. The pooled samples from the “Thiacloprid” and “Flu+Thia” groups had residues of 0.11 mg/kg and 0.20 mg/kg, respectively, in the year 2010-2011 and 0.29 mg/kg and 0.19 mg/kg, respectively, in the year 2011-2012 (Tab. 2).

**Tab. 2.**
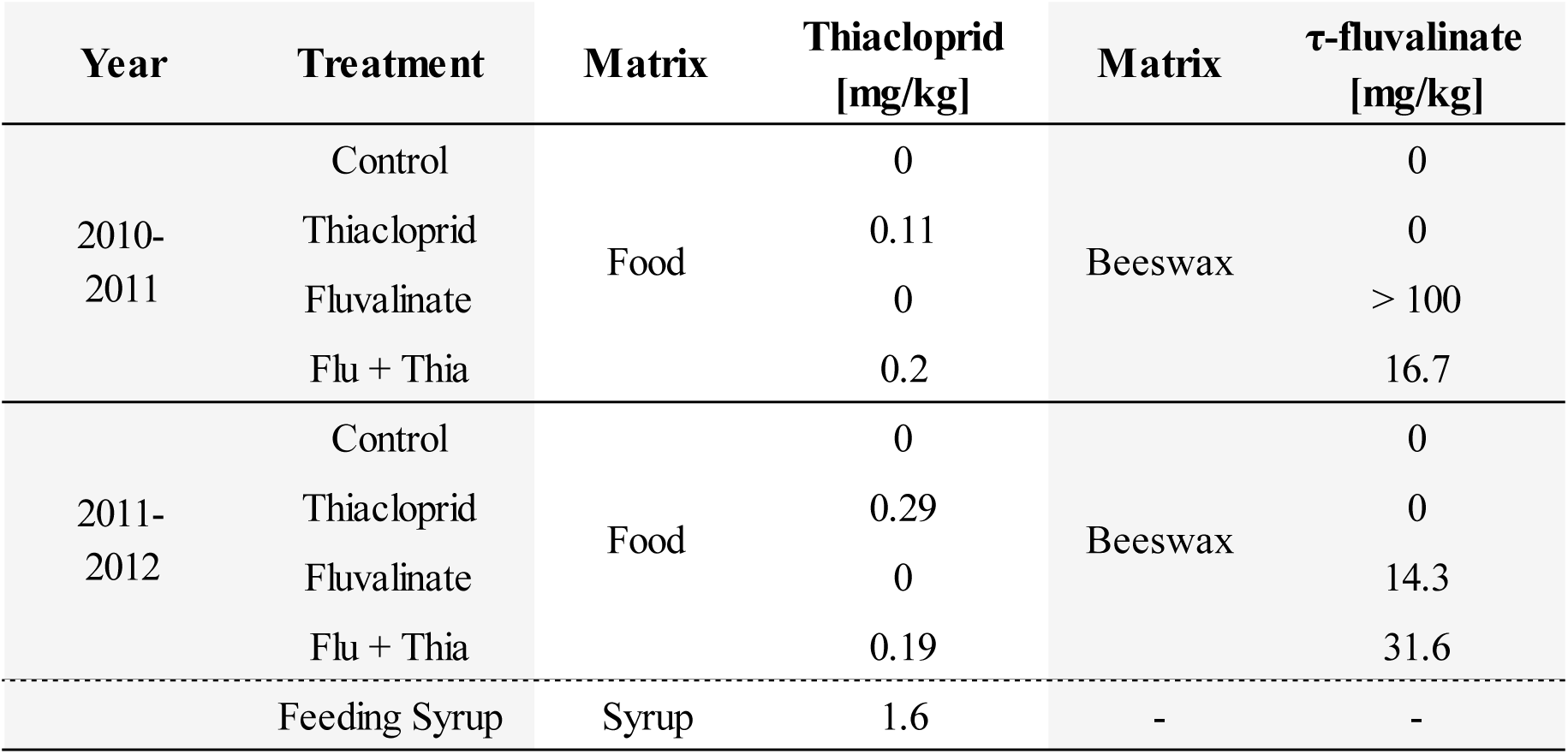
Thiacloprid residues in pooled food (syrup) samples, which was processed by the bees and stored in the honeycombs from all treatment groups in both years (QuEChERS method, LOQ = 0.01 mg/kg). τ-fluvalinate residues in pooled beeswax samples from all treatment groups in both years (SPE & GC-ECD, LOQ = 0.5 mg/kg).

### 3.4 τ-fluvalinate residues

Beeswax was analyzed for τ-fluvalinate residues in both years by solid-phase extraction (SPE) and GC-ECD (LOQ = 0.5 mg/kg). For the analysis, samples from all colonies and the respective groups per year were pooled. All groups without τ-fluvalinate treatment did not have measurable residues in both years. Pooled samples from the “Fluvalinate” and “Flu+Thia” groups had residues of > 100 mg/kg and 16.7 mg/kg, respectively, in the year 2010-2011 and 14.3 mg/kg and 31.6 mg/kg, respectively, in the year 2011-2012 (Tab. 2).

### 3.5 *Varroa* winter treatment

In both years, the winter treatment with oxalic acid killed considerably fewer mites in those groups that have been continuously treated with the acaricide τ-fluvalinate (Fig. 3). In the “Control” and “Thiacloprid” groups between 217 to 409 mites were killed through this winter treatment, on average. In 2010, only one single mite was found in the eight τ-fluvalinate treated colonies! However, in both τ-fluvalinate treated groups the number of mites killed by the winter treatment increased in the second year to an average of 15 mites for the “Fluvalinate” group and 68 mites for the “Flu+Thia” group, respectively.

**Fig. 3.**
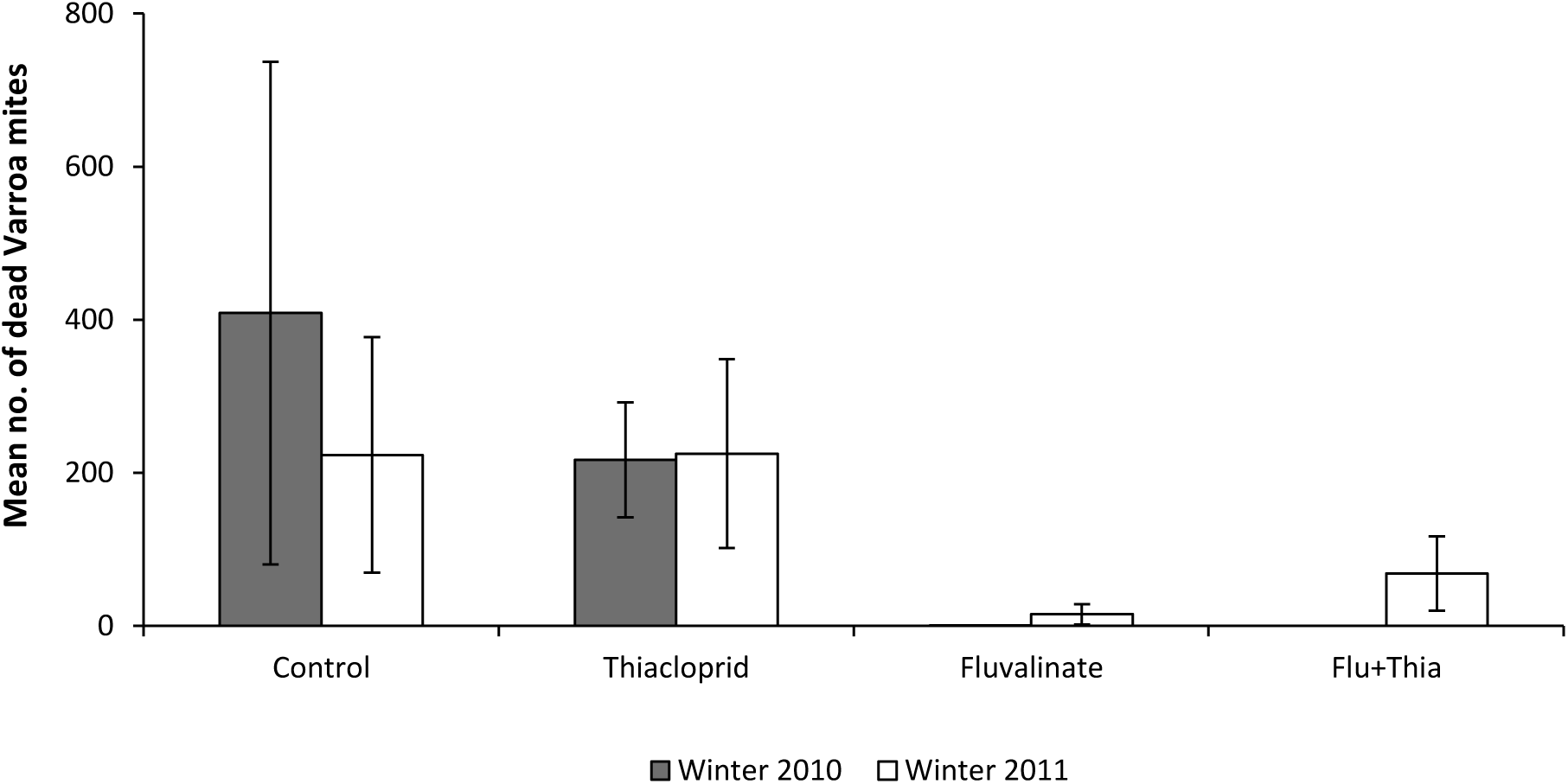
Graph of the dropped *Varroa* mites approximately one week after oxalic acid treatment during winter time (2010 and 2011). In both years a considerably lower number of dead mites could be detected in the τ-fluvalinate treated vs. the untreated groups.

## 4 DISCUSSION

We here analyzed the effects of two commonly used pesticides on the population dynamics and the overwintering success of free flying honey bee colonies. The pesticides belong to two different substance classes, one a neonicotinoid insecticide and the other a synthetic pyrethroid widely used as acaricide to combat varroa mites. For both, the insecticide and the acaricide, the applied dosages represent worst case scenarios. Thiacloprid is meanwhile frequently found as residue in pollen and honey, presumably due to the application in flowering oilseed rape and fruit production. Maximum peak concentrations of thiacloprid in bee products such as nectar, honey or pollen range from ~0.05 to 1 mg/kg across the globe (EFSA, 2016; Genersch et al., 2010; Laaniste et al., 2016; Mitchell et al., 2017; Mullin et al., 2012; Pohorecka et al., 2012; Smodis Skerl et al., 2009) but rarely exceed the average level of 0.2 mg/kg (reports of the German Bee Monitoring, see Rosenkranz et al., 2016). It should be mentioned that 0.2 mg/kg is also the maximum value for thiacloprid residues accepted for honey in the EU (EFSA, 2016). The continuous long-term feeding of 1.6 mg/kg thiacloprid to our experimental colonies resulted indeed in residue levels of this magnitude ranging from about 0.1 to 0.3 mg/kg in the stored food. It is interesting to note the significant 8-fold-decrease from the concentration in the original feeding syrup to the honey bee processed syrup stored in the honeycombs. This decrease might be due to a dilution effect, as all colonies could forage and had access to various nectar sources. Furthermore, Iwasa et al. (2004) and Brunet et al. (2005) reported that cyano-substituted neonicotinoids such as thiacloprid and acetamiprid appear to be metabolized more quickly by the honey bee compared to nitro-substituted ones (i.e. imidacloprid, clothianidin). The enzyme that metabolizes thiacloprid very efficiently but lacking impact against imidacloprid was recently identified as a single cytochrome P450, CYP9Q3 (Manjon et al., 2018). As we did not analyze metabolites, this could additionally have contributed to decrease the in-hive concentration of the pesticide by bees processing the syrup.

For τ-fluvalinate, likewise high maximum residue values are reported. Due to their lipophilic property residues are concentrated and accumulated within the beeswax and can exceed 15 mg/kg (Berry et al., 2013) which is in the range of τ-fluvalinate residues in our experimental colonies after long-term treatment with Apistan® strips. Bogdanov et al. (1998) confirmed an increase of residues with the duration of the strip exposition with a plateau of about 40 to 60 mg/kg after six months whereas other authors found values between 6.6 and 200 mg/kg (Mullin et al., 2010; Adamczyk et al., 2010; Tsigouri et al., 2004).

However, even these residue levels of thiacloprid and τ-fluvalinate are considered to have no acute toxicity to bees or brood (Iwasa et al., 2004; Sanchez-Bayo and Goka, 2014). In our worst case approach we examined whether a long-term exposure to field-realistic peak concentrations of the two pesticides - applied alone or in combination - impairs the development of honey bee colonies under field conditions. In two approaches performed in two consecutive years and using an identical experimental setup we could not detect any negative impact of the treatments on the population of bees and brood and on the overwintering of the colonies. Our moderate overwintering losses of about 15 % (20 % in the first and 8 % in the second winter) are within the range of common winter losses in free flying colonies in Germany and United States (Genersch et al., 2010; Lee et al., 2015) and affected all except the “Fluvalinate” group. Probably, the higher mite load in the untreated groups has contributed to these slightly higher overwintering losses. The mite infestation was quantified in late autumn/winter by an oxalic acid treatment which is known to be highly effective against *Varroa* mites, given that bees are in their winter cluster without brood (Rademacher and Harz, 2006). With the treatment we could also verify that the colonies treated with τ-fluvalinate were sufficiently exposed to this compound during the season, resulting in lower dead mite drops compared to the two groups not treated with τ-fluvalinate. Remarkably, in the winter treatment of the second season our colonies already showed signs of an established τ-fluvalinate resistance in the *Varroa* mite population at our apiary. Such resistance was often reported in the past all over the world (Lodesani et al., 1995; Elzen et al., 1999; Gracia-Salinas et al., 2006; Alissandrakis et al., 2017).

In both years the population of bees and brood was evaluated eight times in a total of 8 - 9 colonies per treatment group. Only in very few cases significant group differences were recorded. In the first year (2010/2011), the control colonies were slightly weaker at the start of the experiment in spring/summer but revealed no differences any more in the autumn and after-winter evaluations. Although all experimental colonies were established from artificial swarms of approximately the same weight it is not unusual that there are small differences in the first weeks of development in newly established honey bee colonies (Imdorf et al., 2008). In the second year (2011/2012) the “Thiacloprid” group had a significant lower number of brood cells in August, however without differences in the two consecutive assessments and without significant effects on the adult bee population. More importantly, there were no group differences at all in the assessments before and after overwintering, indicating no effects of the pesticide treatment on this crucial colony performance. In a previous study performed in observation hives we could already confirm that behavioral traits like flight activity, antennation, grooming and trophallaxis are not affected by the chronic exposure to high concentrations (1 mg/kg) of thiacloprid (Retschnig et al., 2015). The authors therefore assumed a rather weak impact of the pesticide treatment.

Our results are also in agreement with a three-year study of Siede et al. (2017) who chronically applied two different thiacloprid concentrations (0.2 mg/kg and 2 mg/kg) and could also not confirm any negative impairment on colony health and winter survival. Interestingly, they also found a significant lower amount of brood cells in colonies fed with the high thiacloprid concentration but equally to our results no effect on the colony strength or overwintering was noticed. In contrast to other neonicotinoids (Blacquiere et al., 2012) there has been no prove of acute toxicity of thiacloprid to brood; however, according to our results and those of Siede at al. (2017) this aspect should be considered in future approaches. Berry et al., (2013) could also show for τ-fluvalinate, that exposure to high concentrations in beeswax did not have measurable effects on the amount of brood, amount of honey, foraging rate, time required for marked bees released to return to their hive, percentage of released bees that return to the hive, and colony *Nosema* spore loads. In addition, we here could prove for the first time that a combination of this acaricide with the neonicotinoid insecticide did not have measurable synergistic effects at the colony level.

However, our study is in contrast to many laboratory and semi-field studies providing evidence for negative effects of thiacloprid such as elevated mortality under stress (Doublet et al., 2015) or in combination with pathogens (Vidau et al., 2011), impaired navigation (Fischer et al., 2014), reduced immunocompetence (Brandt et al., 2016), disrupted learning and memory functions (Tison et al., 2017) as well as affected social behavior (Forfert and Moritz 2017; Tison et al., 2016). In most of these studies individual bees were exposed to different concentrations of thiacloprid over a certain time period and subsequently challenged to various physiological tests. The findings were then extrapolated to the colony level without confirmation under field conditions. For example, Tison et al. (2016) found foraging behavior and social communication impaired when applying a concentration of 4.5 mg/kg thiacloprid over one week in a free flying feeder experiment. This exposure corresponds to a 23-fold higher concentration than the maximum value for thiacloprid residues accepted for honey in the EU (0.2 mg/kg; EFSA, 2016). It seems unlikely that honey bees are chronically exposed to such high concentrations under realistic field conditions. Additionally, it makes a difference whether pesticides are applied to individual bees under artificial conditions or to bees within a free flying colony. Obviously, the damage threshold of the honey bee colony as a huge social entity is different from the threshold calculated from the effects on individual bees. This “buffering effect” of the colony has frequently been discussed, however without a final explanation of the underlying mechanisms (Straub et al., 2015; Sponsler and Johnson, 2017). Recently, Odemer at al. (2018) could demonstrate that even the highly bee toxic neonicotinoid clothianidin is significantly less toxic when applied to bees that are kept within the social environment of a colony.

Our results might contribute to the current discussion about the ban of neonicotinoids in agricultural practice which recently led to an assessment of the EFSA considering three neonicotinoids (clothianidin, thiametoxam and imidacloprid) a “risk to bees” (EFSA, 2018). It is an important issue for the agricultural production and for environmental protection, whether neonicotinoids with substantially lower bee toxicity should also be banned. Our results indicate that at least for honey bees the risk is low. It is likely that wild bees or other pollinating insects are more susceptible to thiacloprid as it has been shown already for bumble bees (Ellis et al., 2017), however more field data on the population level of wild pollinators are necessary for a reliable risk assessment of thiacloprid.

## ACKNOWLEDGEMENTS

We appreciate the support of the LAB staff for helping with the artificial swarms and colony assessments. This work was supported by the 7^th^ Framework EU Programme Grant “BEE DOC” under Grant number 244956 CP-FP.

